# Adaptation of redox metabolism in drug-tolerant persister cells is a vulnerability to prevent relapse in pancreatic cancer

**DOI:** 10.1101/2022.12.28.522091

**Authors:** Nadine Abdel Hadi, Gabriela Reyes-Castellanos, Tristan Gicquel, Scarlett Gallardo-Arriaga, Emeline Boet, Jean-Emmanuel Sarry, Rawand Masoud, Juan Iovanna, Alice Carrier

## Abstract

Pancreatic Ductal Adenocarcinoma (PDAC) remains a major unresolved disease because of its remarkable therapeutic resistance. Even patients who respond to initial therapy experience relapse in most cases. The mechanisms underlying therapy-acquired resistance supporting relapse are poorly understood. In this study, we aimed to determine the metabolic features of PDAC during relapse, specifically adaptations of mitochondrial and redox metabolism. We used preclinical PDAC mouse models (patient-derived xenografts and murine syngeneic allografts) that present complete regression under initial chemotherapeutic treatment but relapse after a certain time. Relapsed tumors were analyzed *ex vivo* by flow cytometry to measure mitochondrial and redox characteristics. Molecular mechanisms were investigated by quantification of ATP and antioxidants levels, RT-qPCR and bulk RNA-sequencing. Our findings show that mitochondrial metabolism is reprogrammed during relapse, with increased mitochondrial mass, ATP levels, mitochondrial superoxide anions, and total ROS levels, in relapsed compared to control tumors in both models; mitochondrial membrane potential is increased in the xenografts model only. This mitochondrial metabolic reprogramming occurs during treatment-induced regression and at relapse onset. At the molecular level, antioxidant defenses are increased in relapsed tumors and during treatment. These data suggest that treatment-induced oxidative stress may cause the appearance of treatment-adapted cells, known as drug-tolerant persister (DTP) cells. Finally, the combined treatment of arsenic trioxide (ROS inducer) and buthionine sulfoximine (glutathione synthesis inhibitor) is able to completely prevent relapse in PDAC xenografts. In conclusion, targeting redox metabolism via ROS production and antioxidant inhibition is a very promising approach to prevent relapse in PDAC patients.

**Significance:** Mitochondrial and redox metabolisms are reprogrammed during treatment-acquired resistance in pancreatic cancer promoting the survival of drug-tolerant persister cancer cells, opening up new avenues for better therapeutic management of patients.

## INTRODUCTION

Pancreatic Ductal Adenocarcinoma (PDAC) is the most common cancer of the pancreas. To date, pancreatic cancer is the third leading cause of cancer-related deaths in the Western world and is predicted to become the second in the United States by 2026 (1,2). PDAC remains an incurable disease, with the lowest 5-year survival rate after diagnosis of only 11% (2). This situation reflects the slow progress against this disease due to a lack of early diagnosis and major treatment advances (3,4). The poor prognosis of PDAC is linked to a high percentage of locally advanced or metastatic diseases at the time of diagnosis, and the limited effectiveness of antitumor treatments, including immunotherapies (5–7). A small proportion of patients (less than 20%) show a localized tumor eligible for resection at diagnosis. Surgery followed by chemotherapy increases the median survival from 6 to 21 months, but most of the patients experience treatment resistance and develop disease recurrence at some point during treatment or after remission (8,9). It is therefore imperative to identify more effective therapies against this dreadful disease and to better understand the mechanisms of therapy resistance. In addition, to support prevention programs, a better knowledge of the risk factors for developing PDAC is also required, together with the conditions favoring cancer relapse.

Oncologists are confronted with two types of resistance to therapy: primary resistance is an intrinsic non-response to conventional treatments, while acquired resistance is therapy-induced resistance driving relapse (recurrence) after an initial response to treatment (10). The common mechanisms involved in primary chemoresistance include an alteration of drug metabolism, changes in drug targets, apoptosis suppression, activation of intracellular survival signaling, enhanced DNA damage repair, epithelial-mesenchymal transition (EMT), immune escape of cancer stem cells, epigenetic alterations, and aberrant metabolism (10,11). Regarding acquired resistance, a major role has recently been suggested for a subpopulation of drug-tolerant persister (DTP) cancer cells, which survive anti-cancer agent’s treatment through different strategies, and are able to resume proliferation thus from which relapsed tumors emerge (12–14). A common feature of many DTP cancer cells is the adaptation of their cellular metabolism, particularly increased mitochondrial oxidative metabolism, which has been demonstrated in different cancer types (15–20). Another metabolic characteristic of these cells is the overproduction of reactive oxygen species (ROS) and the increase of antioxidant defenses to counteract them, thus allowing cell survival. Such a process includes the increase in glutathione levels, which is commonly observed in various DTP cancer cells (21–23). In addition, phospholipid glutathione peroxidase 4 (GPX4) activity has been shown to increase in DTP cells from melanoma and breast, lung, and colorectal cancer (22,24,25).

In PDAC, the importance of mitochondria in therapeutic resistance was suggested ten years ago by two pioneer studies (26,27). In our laboratory, we showed heterogeneity between PDAC tumors for the two main energy-producing pathways (glycolysis and mitochondrial oxidative phosphorylation [OXPHOS]), and we demonstrated that the chemosensitivity of high OXPHOS tumors can be increased by inhibiting mitochondrial respiratory complex I with phenformin *in vitro* and *in vivo* (28,29). Moreover, we showed that mitochondrial respiration is dependent on fatty acid oxidation (FAO) in all PDAC tumors, and unveiled a synergy between chemotherapy and perhexiline (known as a FAO inhibitor) cytotoxicity (30). We found that an *in vivo* treatment combining perhexiline with standard chemotherapy gemcitabine was able to induce complete regression in one patient-derived xenograft (PDX) model among three that were gemcitabine-resistant (30). Of note, complete regression was obtained for a fourth PDX model treated with gemcitabine alone. Strikingly, we observed relentless tumor relapse after complete regression in these two PDX, mirroring what happens in patients. These relapsed tumors were shown to respond to a second cycle of treatment (30), suggesting that they emerge from residual DTP cancer cells that survived during the first cycle of treatment.

Based on the above data, in the present study, we explored mitochondrial metabolic adaptations in PDAC tumors during relapse, in particular adaptations in redox metabolism. This study was conducted exclusively *in vivo* using our preclinical mouse models (PDX and syngeneic allografts) that relapse after chemotherapy-induced regression.

## MATERIALS AND METHODS

### Animal models

#### Xenograft mouse models of PDAC

We performed subcutaneous xenografts in immunodeficient mice using two different primary human PDAC cells, as we described in Reyes-Castellanos et al. (30). Recipient mice were 5- to 6-week-old athymic female, Swiss nude mice, SOPF (Specific and Opportunistic Pathogen Free) health status, strain Crl:Nu(lco)-Foxn1^nu^ (Charles River, France). To obtain the xenografts, the subcutaneous tumor from an initial mouse donor was removed and finely minced with a scalpel. Then, 150 mg of tumor’s pieces were mixed with 50 μL of Matrigel and implanted with a trocar (10 Gauge) in the subcutaneous space of recipient isoflurane-anesthetized mice. Tumor volume was measured twice per week using a digital caliper, and tumor volume was calculated using the formula V = length x (width)^2^/2.

When xenografts reached ∼ 200 mm^3^ volume, mice were randomly assigned in a treatment group in which the average of all tumors was 200 mm^3^. Treatments were administered by intraperitoneal (IP) injection during one month as follows: gemcitabine 120 mg/kg IP twice a week, or gemcitabine plus perhexiline (120 mg/kg IP twice a week and 5 mg/kg IP every other day, respectively). Vehicle-injected mice (controls) were injected with PBS in the case of gemcitabine controls or 3% DMSO in PBS for combination treatment controls. Mice whose tumor volume reached 1.5 cm^3^ were ethically sacrificed and tumors removed.

The *in vivo* therapeutic effect of arsenic trioxide (ATO, AS_2_O_3_) and L-buthionine sulfoximine (BSO) was evaluated in the xenograft mouse model. Mice were divided into different groups (at least n=4 per condition): PBS (vehicle control), AS_2_O_3_ (0.2 mg/kg), and BSO (0.3 mg/kg). AS_2_O_3_ was administrated IP daily (5 days a week), starting right after the end of the one-month chemotherapy treatment inducing complete tumor regression, and during one, two or three months. BSO was administrated IP 3 times per week for 2 months; BSO treatment started 9 and 16 days (PDAC032T and PDAC084T xenografts, respectively) after the end of the one-month chemotherapy treatment.

#### Syngeneic allograft mouse model of PDAC

Orthotopic syngeneic allografts were generated as previously described (28). The murine KPCluc2 cells were cultured in RPMI medium supplemented with 10% FBS and 600 μg/ml Hygromycine B (ThermoFisher Scientific) for selection of cells containing the luciferase-encoding vector, at 37°C with 5% CO_2_ in a humidified atmosphere. One million of KPCluc2 cells was IP injected into 5- to 6-week-old C57BL/6 female mice (immunocompetent strain, SOPF health status, Charles River, France). Tumoral growth was followed by bioluminescence upon injection of 3 mg luciferin-EF (Promega) using a Photon Imager device (Biospace Lab). Twelve days post-grafting, tumor-bearing mice were randomly assigned to two treatment cohorts (at least n = 5 per condition), vehicle control (treated with PBS) and gemcitabine 120 mg/kg IP twice a week during one month. Mice were sacrificed when they reached the ethical limit point.

All mice were kept under specific pathogen-free conditions and according to the current European regulation; the experimental protocol was approved by the Institutional Animal Care and Use Committee (#16711).

### Tumor dissociation

Xenograft and allograft tumors were dissociated using the gentle MACS™ Octo Dissociator with Heaters and the Tumor Dissociation Kit, mouse (130-096-730), as per manufacturer’s instructions. Briefly, tumors were cut into 2-4 mm^3^ pieces and resuspended in RPMI 1640 media supplemented with 2% FBS (Thermo Scientific, Waltham, MA, USA). These pieces then underwent mechanical and enzymatic digestion for 1 hour. Immediately following dissociation, all single-cell suspensions were filtered using a MACS SmartStrainer (70 μm). The suspensions were centrifuged for 7 min at 300 g at room temperature, the supernatant aspirated, and the cells were resuspended in RPMI 2% FBS media. Cells were counted by flow cytometry using MACSQuant-VYB (Miltenyi Biotec).

### Flow cytometry *ex vivo*

#### Mitochondrial mass and mitochondrial membrane potential measurements

Mitochondrial mass and mitochondrial membrane potential measurements were performed using MitoTracker Deep Red (M22426), and the MitoProbe™ TMRM Kit (M20036), respectively. Briefly, 200,000 cells were collected after tumor dissociation, centrifuged and labeled with MitoTracker and TMRM to a final concentration of 200 nM and 20 nM in PBS at 37°C for 10 min and 30 min, respectively. Then, 10,000 events per sample were acquired in a MACSQuant-VYB (Miltenyi Biotec), and data analysis was performed using the FlowJo software.

#### Total ROS and mitochondrial superoxide anions detection

Total ROS and mitochondrial superoxide anions measurements were performed using CellRox Orange (C-10443), and MitoSOX Red (M36008), respectively. Briefly, after tumor dissociation, 200,000 cells were collected, centrifuged and labeled with CellROX and MitoSOX to a final concentration of 5μM and 10 μM (in PBS for MitoSOX, and culture medium for CellRox) for 30 min and 20 min at 37°C, respectively. After incubation, cells were centrifuged, and resuspended in PBS 1x for flow cytometry analysis. 10,000 events per sample were acquired in a MACSQuant-VYB (Miltenyi Biotec), and data analysis was performed using the FlowJo software.

### ATP level measurement

Total ATP level was measured using the Cell viability assay (Cell-Titer Glo Kit; Promega) according to the manufacturer’s instructions. Briefly, 50,000 cells were resuspended in 100 μL of medium and distributed in four replicates in a 96-well flat-bottom plate. Cells were then treated with PBS (Control), 1μM oligomycin (inhibitor of mitochondrial respiration), and 100mM 2-DG (inhibitor of glycolysis). Following one hour of incubation at 37°C, 100 μL of Cell Titer Glo reaction mix solution was added to each well for a final volume of 200 μL. Plates were then analyzed by luminescence using Tristar LB 941 apparatus (Berthold technologies). The background relative light unit (RLU) was subtracted from each RLU value. By comparing the different conditions, total ATP (PBS condition) and percentages of both mitochondrial and glycolytic ATP were determined: mitochondrial ATP = Total ATP - ATP (oligomycin); mitochondrial ATP (%) = mitochondrial ATP / Total ATP *100; glycolytic ATP = Total ATP - ATP (2DG); glycolytic ATP (%) = glycolytic ATP / Total ATP *100.

### Quantification of small antioxidant molecules

#### GSH/GSSG measurement

GSH/GSSG-Glo Assay kit (Promega, V6611) was used following manufacturer’s protocol with some modifications. Briefly, 10 mg of tumor tissue was crushed using Precellys® Evolution, resuspended in 50 μL of PBS/EDTA, and distributed in a 96-well plate. After homogenization, 50μl of Total Glutathione Lysis Reagent (for Total glutathione measurement) and Oxidized Glutathione Lysis Reagent (for GSSG measurement) were added. Luciferin Generation Reagent and Detection Reagent were added to all wells and luminescence was recorded using Tristar LB 941 apparatus (Berthold technologies). By comparing the different conditions, GSH/GSSG ratio was calculated using the following equation: GSH/GSSG = (total glutathione RLU - GSSG RLU) / (GSSG RLU/2).

#### NADPH measurement

10 mg of tumor tissue were crushed using Precellys® Evolution, resuspended in 50 μL of PBS/EDTA, and distributed in a 96-well plate. Measurement was performed according to the manufacturer’s protocol. Briefly, 50 μl of NADP/NADPH-Glo™ Detection Reagent (Promega, G9081) was added to each well and incubated for 30 minutes at RT. Luminescence was recorded using Tristar LB 941 apparatus (Berthold technologies).

### Gene expression analysis by RT-qPCR

Total RNA was isolated from 20-25 mg of tumor piece using both TRIzol (Invitrogen) and the Qiagen total RNA isolation kit (Qiagen, ref 74004) to avoid protein and extracellular matrix accumulation in the columns. The tumor piece was lysed in 600 μL RLT Buffer (from Qiagen kit) with 1% β-Mercaptoethanol using beads tube from Precellys lysing kit for hard tissue (ref P000917-LYSK0-A, 3x Cycle 1500 rpm, 15 sec of mix, 10 sec of rest) and directly centrifuged 3 min at 10.000g at RT. Supernatant was transferred in 400μL of TRIzol and incubated for 5 min at RT followed by addition of 150μL of chloroforme. Tubes were carefully mixed and incubated 3 min at RT before centrifugation (12.000g, 5min, 4°C). The transparent upper phase containing RNAs was transferred in a tube containing 500 μL of 70% ethanol, mixed and transferred inside Qiagen RNA isolation kit columns according to manufacturer’s instructions which were followed until the end of extraction. RNA samples were subjected to reverse-transcription (RT) using the Go Script reagent (Promega) following manufacturer’s instructions. Next, Real-Time quantitative PCR was performed in triplicates using Takara reagents and the Stratagene cycler Mx3005P QPCR System. Raw values were normalized with the housekeeping gene TBP1 for the same cDNA sample. We used the human primers for Nrf2, HO-1, SLC7a11, and GPX4 involved in the antioxidant defense, and PGC-1α and TFAM involved in mitochondrial metabolism. Primer sequences can be found in (31). For each RNA sample, RT reaction was done twice to generate 2 batches of cDNA, which were amplified by qPCR in three independent experiments for each gene.

### Transcriptomic analysis by bulk RNA sequencing

#### Total RNA isolation from tumors

RNA was extracted from PDAC xenografts using the RNeasy Mini kit (Qiagen) as described above. RNA integrity and concentration were assessed using the Agilent 2100 Bioanalyzer (Agilent Technologies, Palo Alto, CA). The average RIN (RNA integrity number) values for all samples was comprised between 9.3 and 10, ensuring a high quality of isolated RNAs.

#### Bulk RNA sequencing

The preparation of mRNA libraries was realized following manufacturer’s recommendations (kapa mRNA HyperPrep from ROCHE). Final samples pooled library prep were sequenced on ILLUMINA Novaseq 6000 with S1-200cycles cartridge (2×1600Millions of 100 bases reads), corresponding to 2×30Millions of reads per sample after demultiplexing.

#### Bulk RNA sequencing data analysis

Quality control has been performed on the fastq files using FastQC (v0.11.9) (http://www.bioinformatics.babraham.ac.uk/projects/fastqc). To map the sequenced reads to the human reference genome, we made use of STAR (v2.7.3a). From these mapped reads, gene counts were then quantified using featureCounts (v2.0.1). Starting from the raw gene counts, normalization and differential expression analysis have then been performed using DESeq2 (v 1.22.2).

#### Gene Set Enrichment Analysis (GSEA)

Gene set enrichment analysis (GSEA) was performed using GSEA v4.1 tool (Broad Institute). Following parameters were used: Number of permutations = 1000, permutation type = gene set, Chip = Human_Ensembl_Gene_ID_MSigDB.v2022.1.Hs. chip. Other parameters were left at default values. The normalized enrichment scores (NES) were computed from PDAC032T relapse group compared to PDAC032T control group. Genes with FDR q-value < 0.05 and fold change >= 1.5 were considered of interest.

### In vitro assays

#### Cell viability

The human PDAC032T and PDAC084T primary PDAC cells were cultured in serum-free ductal media (SFDM) at 37°C with 5% CO_2_ in a humidified atmosphere as reported previously (28,30). SFDM is a complex medium supporting the PDAC primary cell growth and containing DMEM-F12, nicotinamide, glucose, hormones, growth factors and Nu-serum providing a low-protein alternative to fetal bovine serum (FBS). Cells were seeded in 96-well plates in triplicates (5,000 cells per well) and the corresponding treatment was administered the day after. Cells were treated with perhexiline (7 μM), gemcitabine (1 μM), or the combination for 24 hours. These treatments were done in the presence and absence of NAC (2.5 mM). Next, cell viability was determined by the Crystal violet viability assay, which is independent of cell metabolism. For this, cells were fixed in glutaraldehyde (1%), washed twice with PBS, stained with Crystal violet (0.1%) for 10 min, and then washed three times with PBS. Crystals were solubilized in SDS (1%), and absorbance was measured at 600 nm using an Epoch-Biotek spectrophotometer.

#### Total ROS measurement by flow cytometry

Cells were seeded in 12-well plates in duplicates (200,000 cells/1 ml media/well) and the day after, treatments were administered. Cells were treated with perhexiline (7 μM), gemcitabine (1 μM) or the combination for 24 hours. After treatments, the media was supplemented with CellROX Orange at a final concentration of 5 μM. Cells were incubated for 30 minutes at 37°C, then harvested with pre-warmed Accutase (Gibco) and resuspended in PBS for flow cytometry analysis. 10,000 events per sample were acquired in a MACSQuant-VYB (Miltenyi Biotec), and data analysis was done with the FlowJo software.

### Statistical analysis

Results are expressed as the mean ± SEM or SD of duplicates or triplicates, and at least two or three independent experiments were done for each analysis. Statistical analysis of data was performed with GraphPad Prism 8 (GraphPad Software) using two-tailed unpaired Student’s t-test. *P* values < 0.05 were considered to be statistically significant (\**P* < 0.05, \*\**P* < 0.01, \*\*\**P* < 0.001, and \*\*\*\**P* < 0.0001).

### Data availability

The data generated in this study will be made publicly available by submission in a repository upon publication acceptance.

## RESULTS

### The mitochondrial metabolism is reprogrammed in relapsed PDAC tumors

We used two subcutaneous PDAC xenograft models, PDAC032T and PDAC084T, which always relapse after complete regression induced by the chemotherapy, namely gemcitabine alone or combined with perhexiline (FAO inhibitor), respectively (Fig. 1A and B). We analyzed mitochondrial characteristics and ROS levels in relapsed tumors at the ethical end point. Flow cytometry data presented in Fig. 1C and Supplementary Fig. S1 illustrate that mitochondrial mass, mitochondrial membrane potential (MMP, reflecting respiratory activity), mitochondrial superoxide anion (O_2_^.-^) level, and total ROS level are higher in relapsed tumors (in both models) compared with controls (non-treated). Interestingly, only a portion of the increase in MMP and mitochondrial O_2_^.-^ comes from the increase in mitochondrial mass, as shown in the normalized data (Supplementary Fig. S2A). In addition, we found a higher level of total ATP in the relapsed PDAC084T xenograft, a higher percentage of ATP produced by mitochondria in both relapsed xenografts, and a higher percentage of ATP produced by glycolysis in the relapsed PDAC032T xenograft (Fig. 1D and Supplementary Fig. S2B), suggesting higher energy production in both settings.

**Figure 1.**
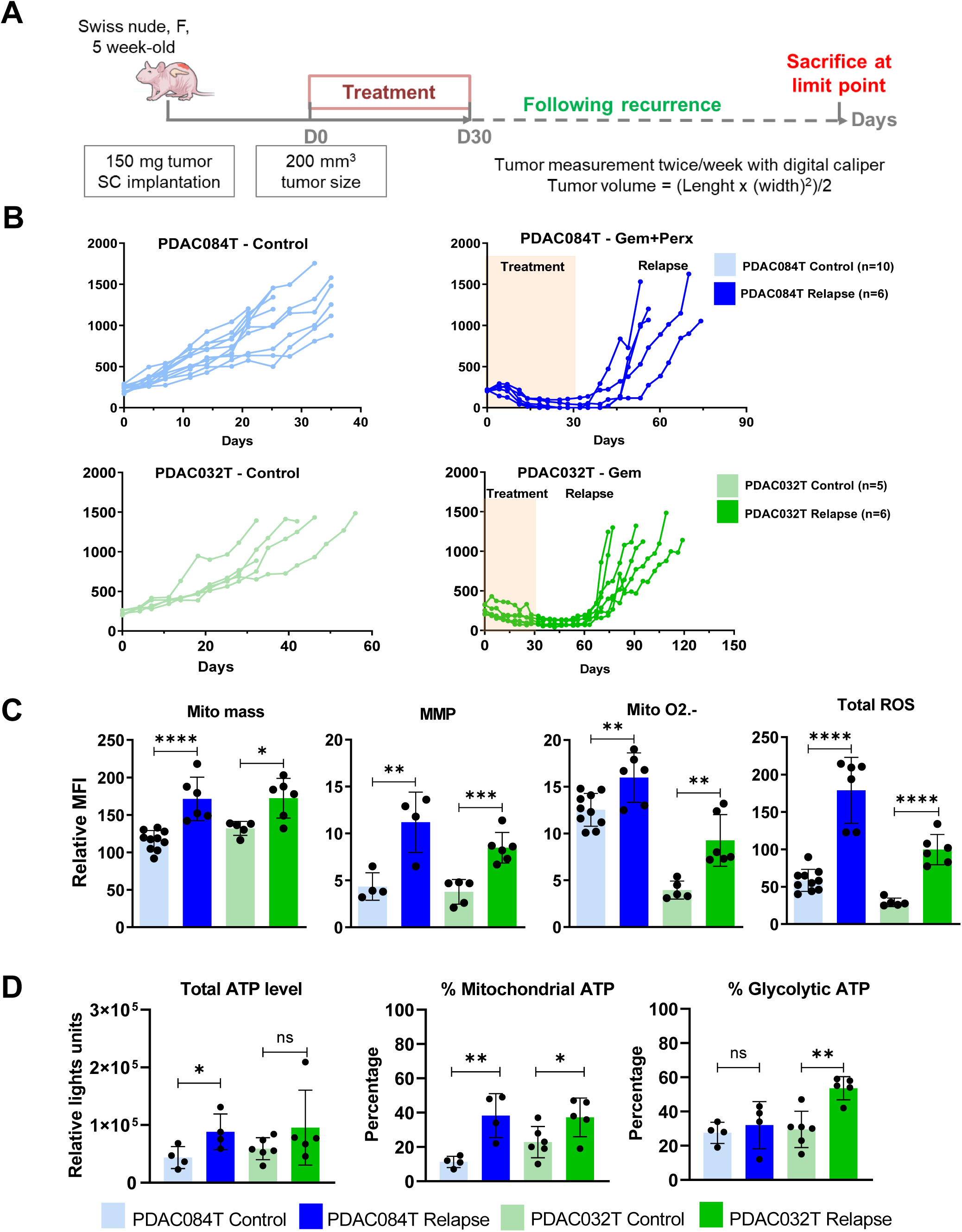
Mitochondrial and redox metabolisms are reprogrammed in relapsed PDAC xenografts. **(A)** Schematic of experimental protocol. Pieces of tumor from PDX were implanted in the subcutaneous space of recipient female Swiss nude mice. When tumors reached 200 mm^3^ volume, mice were assigned to treatment groups and treated for one month. After one month, the treatment was stopped and the recurrence was followed. Mice were sacrificed according to the ethical limit point. **(B)** Tumor volume in two different PDAC xenografts (PDAC084T and PDAC032T) treated during one month with gemcitabine (120 mg/kg IP twice a week), gemcitabine plus perhexiline (120 mg/kg IP twice a week and 5 mg/kg IP every other day, respectively), and Control. PDAC xenografts always relapse after complete regression induced by the chemotherapy, gemcitabine alone or combined with perhexiline for PDAC032T (right) and PDAC084T (left), respectively. **(C)** Mitochondrial mass (Mito mass), mitochondrial membrane potential (MMP), mitochondrial superoxide anions (Mito O2.-) and total reactive oxygen species (Total ROS) levels were measured by flow cytometry with the MitoTracker DeepRed, TMRM, MitoSOX red, and CellROX orange fluorescent probes, respectively, in relapsed tumors at ethical limit point. Median fluorescence intensity (MFI) is shown relative to that of unlabeled cells. **(D)** Total ATP level and percentages of mitochondrial and glycolytic ATP were measured using the Cell viability assay (Cell-Titer Glo Kit). For this, specific inhibitors were used: oligomycin (1μM) and 2-DG (100mM) to determine percentages of mitochondrial and glycolytic ATP, respectively. Unpaired student’s T-test was used for statistical analyzes comparing each group with the untreated group. *, **, *** and **** correspond to p<0.05, 0.01, 0.001, and 0.0001, respectively; ns: non-significant difference.

Metabolic variations during relapse were also observed in an immunocompetent PDAC mouse model, namely syngeneic allografts (Fig. 2) that we reported previously (28). In this model, murine luciferase-expressing PDAC cells implanted into the peritoneal cavity of syngeneic recipient mice grow preferentially in the pancreas, thus providing a very convenient orthotopic allograft model in which tumor growth and regression can be monitored by bioluminescence (Fig. 2B). All tumors respond to gemcitabine treatment and most regress either completely or partially, and all relapse during or after the end of the 1-month treatment (Fig. 2B and C). As for the xenografts presented above, mitochondrial mass, mitochondrial O_2_^.-^ and total ROS levels are higher in relapsed tumors compared to controls (non-treated), but the difference is significant only for tumors that have fully regressed for mitochondrial mass and total ROS level. In contrast to the xenografts, MMP is decreased in relapsed tumors compared to controls, but the difference is not significant for fully regressed tumors. Only part of the increase in mitochondrial O_2_^.-^ comes from the increase in mitochondrial mass, whereas the decrease in MMP is amplified when normalized by mitochondrial mass (Supplementary Fig. S2C). Next, we found a higher level of total ATP in relapsed allografts (Fig. 2E), suggesting higher energy production during relapse.

**Figure 2.**
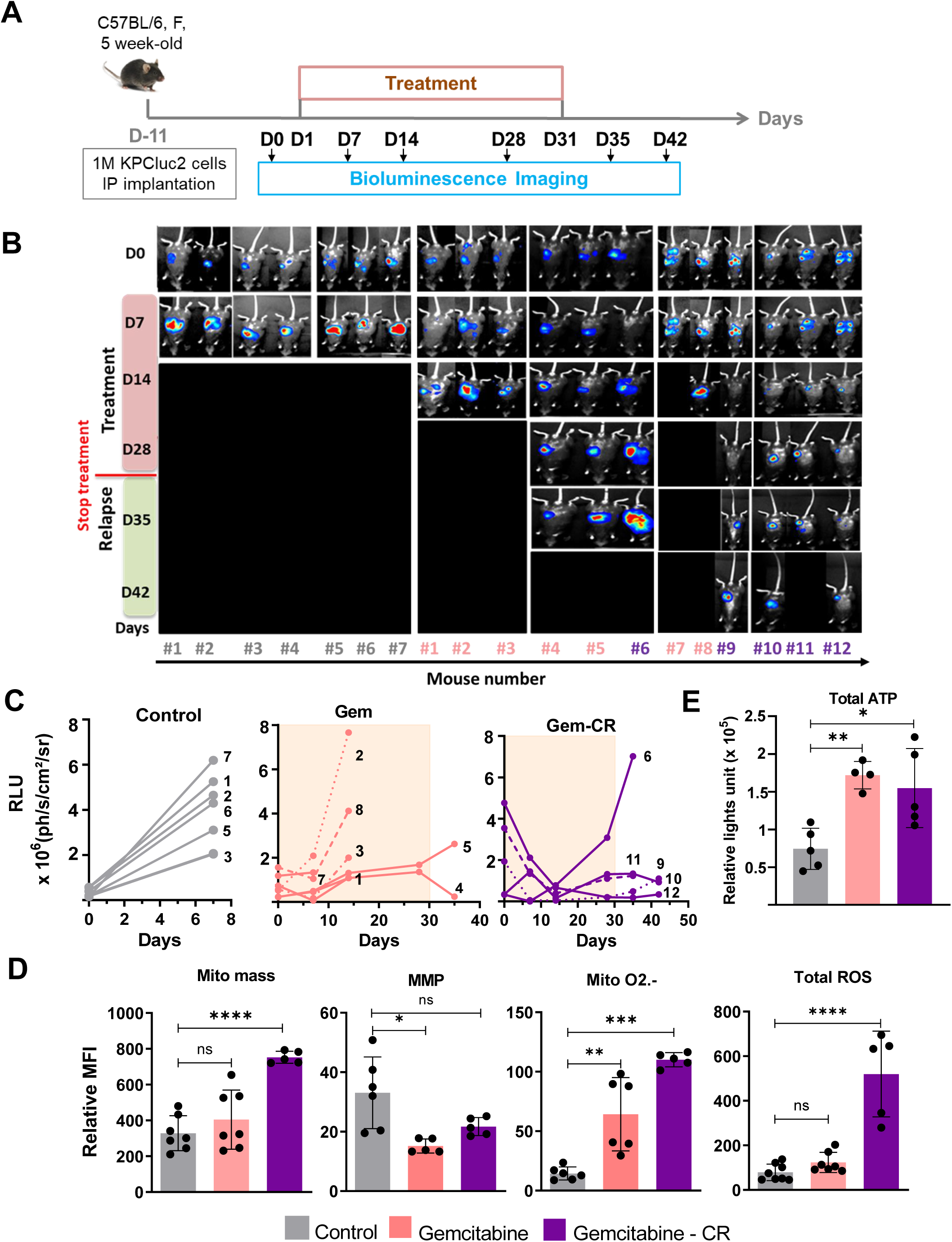
Mitochondrial and redox metabolism are reprogrammed in relapsed PDAC allografts. **(A)** Schematic of experimental protocol. Orthotopic syngeneic allografts were generated by intraperitoneal injection of one million of the murine cells KPC luc2 into 5-week-old female C57BL/6 immunocompetent mice. Tumoral growth was followed by bioluminescence using a Photon Imager device (Biospace Lab). Twelve days post-grafting, tumor-bearing mice were randomized to two treatments cohorts (at least n = 5 per condition), vehicle control (treated with PBS) and gemcitabine 120 mg/kg twice a week by intraperitoneal injection during one month. Mice were sacrificed according to the ethical limit point**. (B-C)** Representative *in vivo* bioluminescence imaging data (**B**) and bioluminescent signal quantification (**C**), at different times after orthotopic implantation of KPCluc2 cells untreated (Control) or treated with gemcitabine. The tumors of mice 1, 2, 3, 4, 5, 7 and 8 respond to treatment with Gem but do not completely regress (Gem group); their number is in pink color. However, those of mice 6, 9, 10, 11, and 12 regressed completely (Gem-CR group); their number is in purple. The bioluminescent signal is expressed in photons per second per square centimeter per steradian (ph/s/cm^2^/sr). **(D)** Mitochondrial mass, MMP, Mitochondrial O2.- and total ROS level were measured in tumors as in Figure 1. Median fluorescence intensity (MFI) is shown relative to unlabeled cells. **(E)** Total ATP level was measured using the cell viability assay (Cell-Titer Glo Kit). Unpaired student’s T-test was used for statistical analyzes comparing each group with that of the untreated. *, **, *** and **** correspond to p<0.05, 0.01, 0.001, and 0.0001, respectively; ns: non-significant difference.

Finally, we performed a transcriptomic study by bulk RNA-sequencing of PDAC032T relapsed tumors compared to non-treated (controls). We applied Gene Set Enrichment Analysis (GSEA) on the RNA-seq data and found that relapsed tumors positively correlate with changes in OXPHOS, MOOTHA_VOXPHOS, electron transport chain (ETC), and the citric acid (TCA) cycle gene signatures (Supplementary Fig. S3A). Furthermore, we found that the expression of genes encoding key transcriptional regulators involved in mitochondrial homeostasis such as Peroxisome Proliferator-activated Receptor–γ Coactivator 1 alpha (*PPARGC1A*) encoding PGC-1α (but not mitochondrial transcription factor A, *TFAM*) was increased in relapsed tumors (Supplementary Fig. S3B).

Collectively, these data show increases in mitochondrial and redox metabolisms in tumors that have relapsed after full treatment-induced regression compared to non-treated tumors, suggesting metabolic reprogramming imposed by the chemotherapy.

### Mitochondrial metabolic reprogramming occurs during treatment-induced regression

We wondered when metabolic changes occurred, in other words, whether they were related to tumor growth during relapse or whether they preexisted relapse. To answer this question, we analyzed xenografts during treatment before complete regression (“under treatment” setting) and at the onset of relapse (“start of relapse” setting), comparing them to untreated tumors of similar size (100-300 mm^3^) (Fig. 3A). We observed that all analyzed parameters were increased in both settings and in both xenografts (Fig. 3B and C, and Supplementary Fig. S4), as observed in relapsed tumors at the end point. Importantly, mitochondrial mass was the parameter that was most increased, explaining the attenuation or even reversal of changes in MMP and mitochondrial superoxide after normalization with mitochondrial mass (Supplementary Fig. 4A and B). These data suggest that the drug treatment, i.e., gemcitabine alone or combined with perhexiline for PDAC032T and PDAC084T xenografts, respectively, could promote the survival of treatment-adapted cells while inducing the death of most tumor cells by oxidative stress associated with the treatment. Indeed, *in vitro* treatments induce an increase in ROS level that can be prevented by supplementation with the antioxidant NAC, and a redox-induced loss of cell viability (Supplementary Fig. S5).

**Figure 3.**
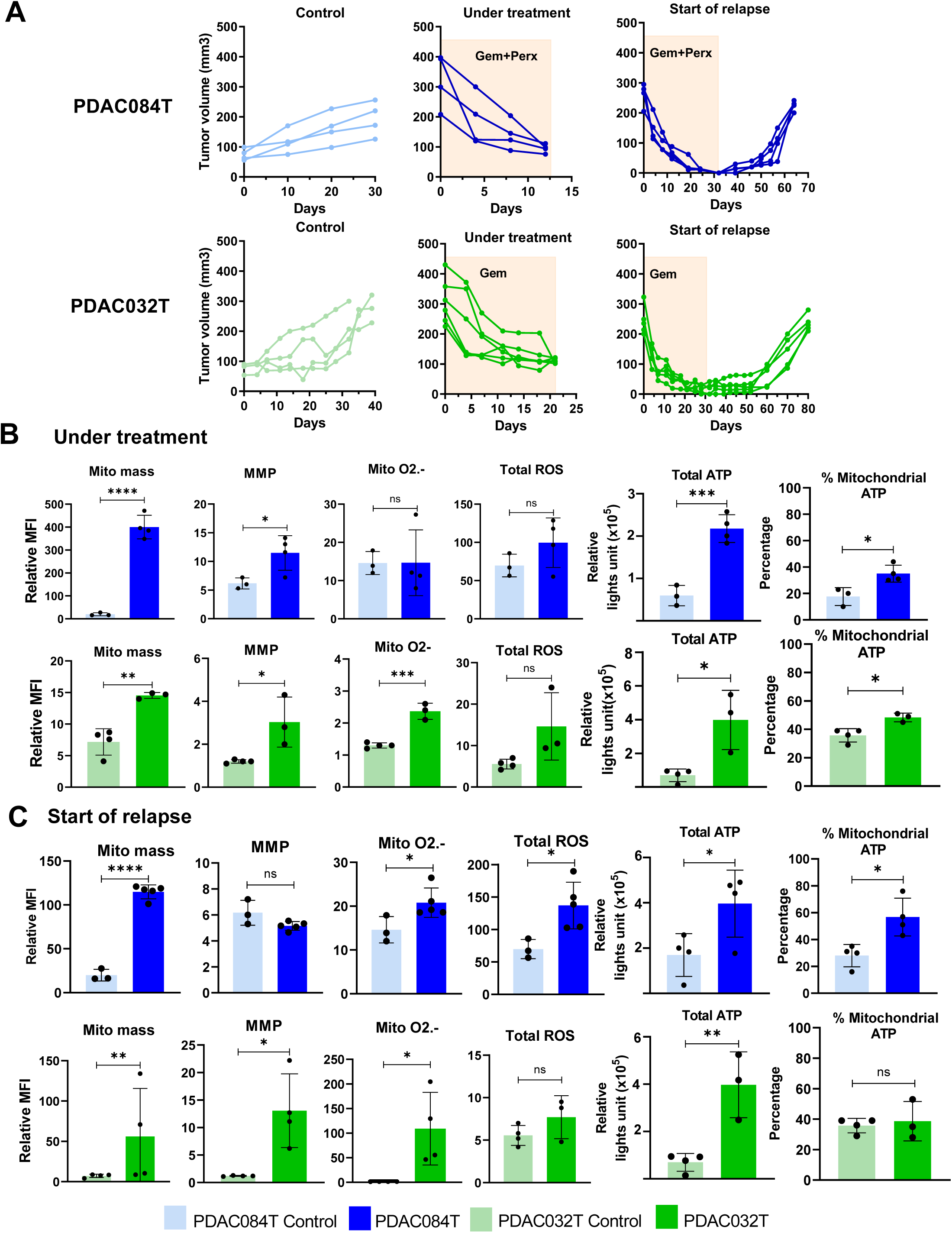
Mitochondrial metabolic reprogramming occurs during treatment-induced regression in PDAC xenografts. **(A)** Tumor volume in PDAC084T (top), and PDAC032T (bottom) during Gem+Perx or Gem treatment alone, respectively, before complete regression (“under treatment” context) and at the onset of relapse (“start of relapse” context), compared to non-treated tumors (Control) of similar size (100-300 mm^3^). For the control groups, we show the measurement of tumor volumes from two weeks after inoculation. For the treated groups, we show the measurement of tumor volumes from the start of treatment. **(B-C)** Mitochondrial mass, MMP, Mitochondrial O2.- and total ROS level were measured in tumors under treatment **(B)** and at the start of relapse **(C).** Total and mitochondrial ATP were measured in both context **(B-C)** using cell viability assay (Cell-Titer Glo Kit). Significance was determined by Student’s T-test. *, **, *** and **** correspond to p<0.05, 0.01, 0.001, and 0.0001, respectively; ns: non-significant difference.

Overall, these data show that chemotherapy treatment affects mitochondrial and redox metabolisms, and suggest that certain cells that survive thanks to metabolic adaptations (residual or persister cells) are responsible for relapse.

### Antioxidant defenses are increased in relapsed tumors and in tumors during treatment

A crucial metabolic adaptation observed in cancer cells during tumor development is the exacerbation of antioxidant defenses to cope with the increased production of ROS, which allows the survival of the adapted cells (32,33). This adaptation is also observed during oxidative stress induced by chemotherapy or radiotherapy (34,35). The main antioxidant defenses are antioxidant enzymes such as glutathione peroxidases (GPXs), superoxide dismutases, and catalase, as well as small antioxidant molecules such as glutathione and NADPH that are found to be abundant in cancer cells (schematic in Fig. 4A). In addition, the nuclear factor erythroid 2-related factor 2 (Nrf2) plays a crucial role in antioxidant gene expression; it can be controlled both at the level of expression and cellular localization since it is translocated to the nucleus only when ROS levels increase. One of the target genes of Nrf2 is *Slc7a11*, which encodes the cystine/glutamate transporter (xCT), which allows the entry of cystine into cells as a precursor of glutathione (tripeptide Glutamate-Cysteine-Glycine). Reduced glutathione (GSH) acts as a co-factor (electron donor) of the GPXs whose antioxidant activity generates oxidized glutathione (GSSG) that is reduced by the glutathione reductase using NADPH as a co-factor.

**Figure 4.**
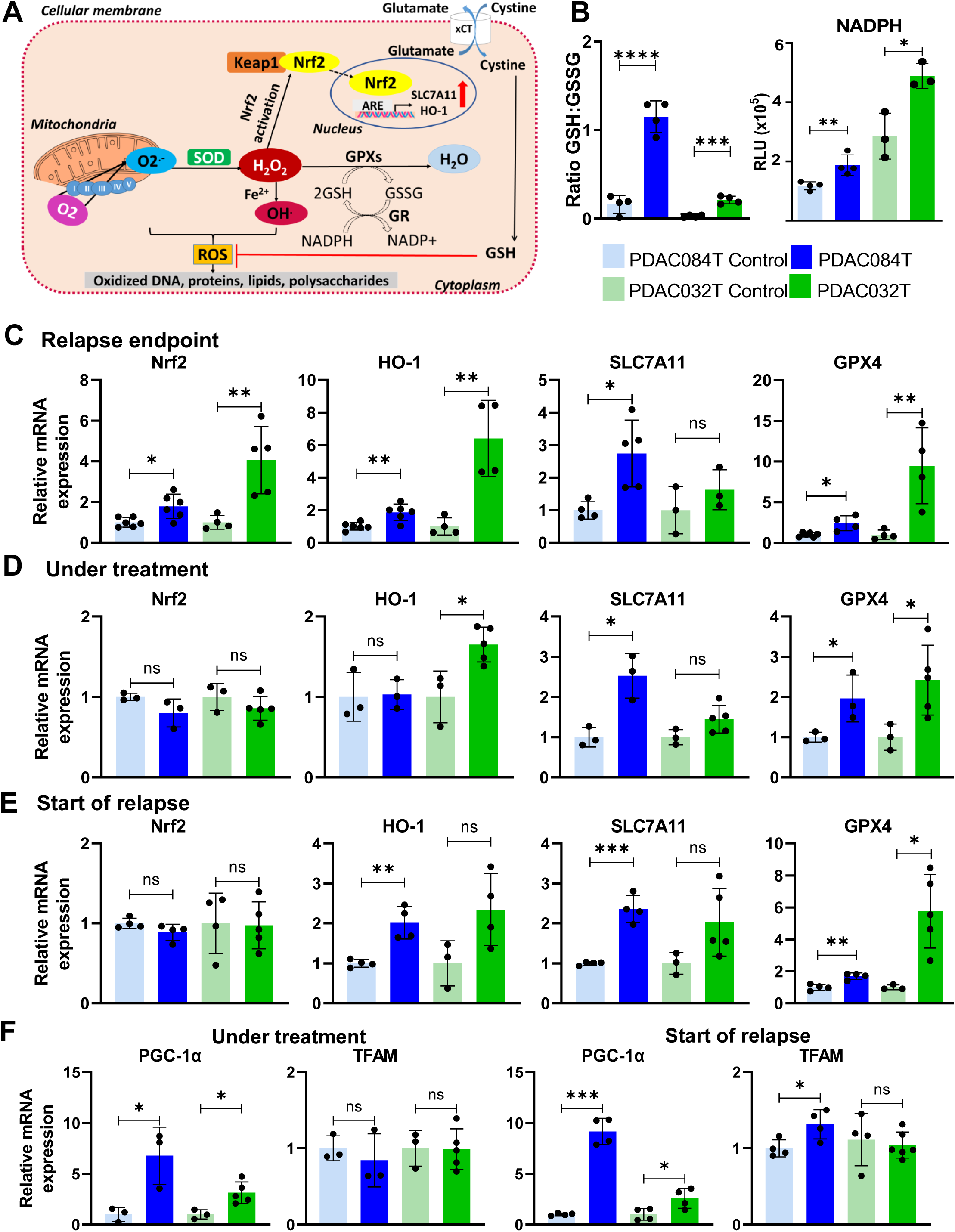
Antioxidant defenses are increased in relapsed tumors and in tumors during treatment. **(A)** Simplified schematic representation of ROS generation and enzymatic antioxidant defenses. Superoxide anion (O_2_^.-^) is mainly produced by the mitochondrial ETC complexes by partial reduction of molecular oxygen (O_2_). Superoxide is dismutated into hydrogen peroxide (H_2_O_2_) by superoxide dismutases (SOD). H_2_O_2_ is converted to water (H_2_O) by glutathione peroxidases (GPX). Via the Fenton reaction with metal ions Fe^2+^ or Cu^+^, H_2_O_2_ is further reduced to highly reactive hydroxyl radical (OH^·^), thereby damaging biological macromolecules such as DNA, lipids, and proteins. H_2_O_2_ is the main player in redox homeostasis, and can induce the activation of the antioxidant transcription factor Nrf2 through dissociation of the Nrf2-KEAP1 complex, phosphorylation of Nrf2, and its nuclear translocation. In the nucleus, Nrf2 promotes transcription of multiple antioxidant genes such as *SLC7A11* and *HO-1* by binding to the antioxidant responsive elements (ARE) in the promoter region of target genes. Via its entry into cells through the glutamate/cystine antiporter xCT encoded by the *SLC7A11* gene, cysteine can enhance GSH production which is a tripeptide glutamate-cysteine-glycine. The antioxidant function of GSH is mediated by two enzymes: GPX and GR. GPX allows the reduction of H_2_O_2_ by the oxidation of GSH (reduced glutathione) to GSSG (oxidized glutathione). The GSSG is subsequently reduced to GSH by GR at the expense of NADPH used as a cofactor. **(B)** Ratio of reduced glutathione (GSH) / oxidized glutathione (GSSG), and NADPH levels were measured in PDAC relapsed tumor. **(C-D-E)** RT-qPCR analysis of antioxidant gene expression: *Nrf2*, *HO-1*, *SLC7A11* and *GPX4*, in relapsed PDAC tumors at endpoint (**C**), under treatment (**D**) and start of relapse (**E**). **(F)** The mRNA levels of PGC-1α and TFAM was monitored by RT-qPCR in relapsed PDAC tumors under treatment and start of relapse, respectively. Data are representative of 3 independent qPCR experiments. Significance was determined by unpaired student’s T-test. *, **, *** and **** correspond to p<0.05, 0.01, 0.001, and 0.0001, respectively; ns: non-significant difference.

We first quantified glutathione and NADPH in xenografts at relapse end point and found increased levels of GSH and NADPH as well as increased GSH/GSSG ratio (Fig. 4B and Supplementary Fig. S6). We then analyzed expression of several antioxidant genes by RT-qPCR. *Nrf2*, its targets *HO-1* and *Slc7a11*, and *GPX4* were found to be overexpressed in the relapse endpoint context (Fig. 4C).

We then measured the expression of these genes in xenografts under treatment and at the onset of relapse (Fig. 4D and E). *Nrf2* was not found to be overexpressed in these settings, unlike its target genes *HO-1* and *Slc7a11* (although some differences did not reach statistical significance), suggesting that Nrf2 is activated even when not overexpressed. The *GPX4* genes was found overexpressed in all conditions.

We also examined the expression of genes encoding proteins involved in mitochondrial homeostasis (PGC-1α and TFAM) and found them overexpressed in most of the conditions analyzed (Fig. 4F), suggesting that increased mitochondrial mass plays a role in mitochondrial ROS overproduction, generating oxidative stress that is counterbalanced by increased antioxidant defenses.

Collectively, these data suggest that the DTP cancer cells responsible for relapse originate from cells that have adapted to drug-induced oxidative stress by increasing antioxidant defenses.

### ROS accumulation is a vulnerability that can be targeted to prevent relapse

The property of cancer cells to have a high ROS content can be used therapeutically by further increasing the level of ROS to reach the threshold for induction of cell death ^29,30^. This was the course of action we took in an attempt to further increase the level of ROS in persister cells to prevent relapse. To this end, we started treatment just after the end of chemotherapy with arsenic trioxide (ATO), which is known to increase ROS production by inhibiting mitochondrial OXPHOS (36). ATO treatment was first performed in PDAC032T xenograft for 1, 2 or 3 months (Fig. 5A). We observed that ATO was able to delay relapse compared with control (relapse of non-ATO-treated mice) in the case of mice treated with ATO for 1 month. Tumor growth started to resume before the end of treatment in the case of mice treated with ATO for 2 months and 3 months. We analyzed mitochondrial characteristics and ROS level in recurrent tumors and found an increase in mitochondrial mass, MMP, and ROS level in ATO-treated recurrent tumors treated for 1 month compared with the non-ATO-treated control mice (Fig. 5B). In contrast, these characteristics decreased in recurrent ATO-treated tumors treated for 2 or 3 months compared with the control group or even the one-month treated group. Finally, mitochondrial MMP and superoxide anions levels decreased in ATO-treated recurrent tumors at all time points when normalized by mitochondrial mass (Supplementary Fig. S7), as well as ATP level (Fig. 5B), suggesting an ATO-induced energy crisis consistent with its role in inhibiting mitochondrial respiration.

**Figure 5.**
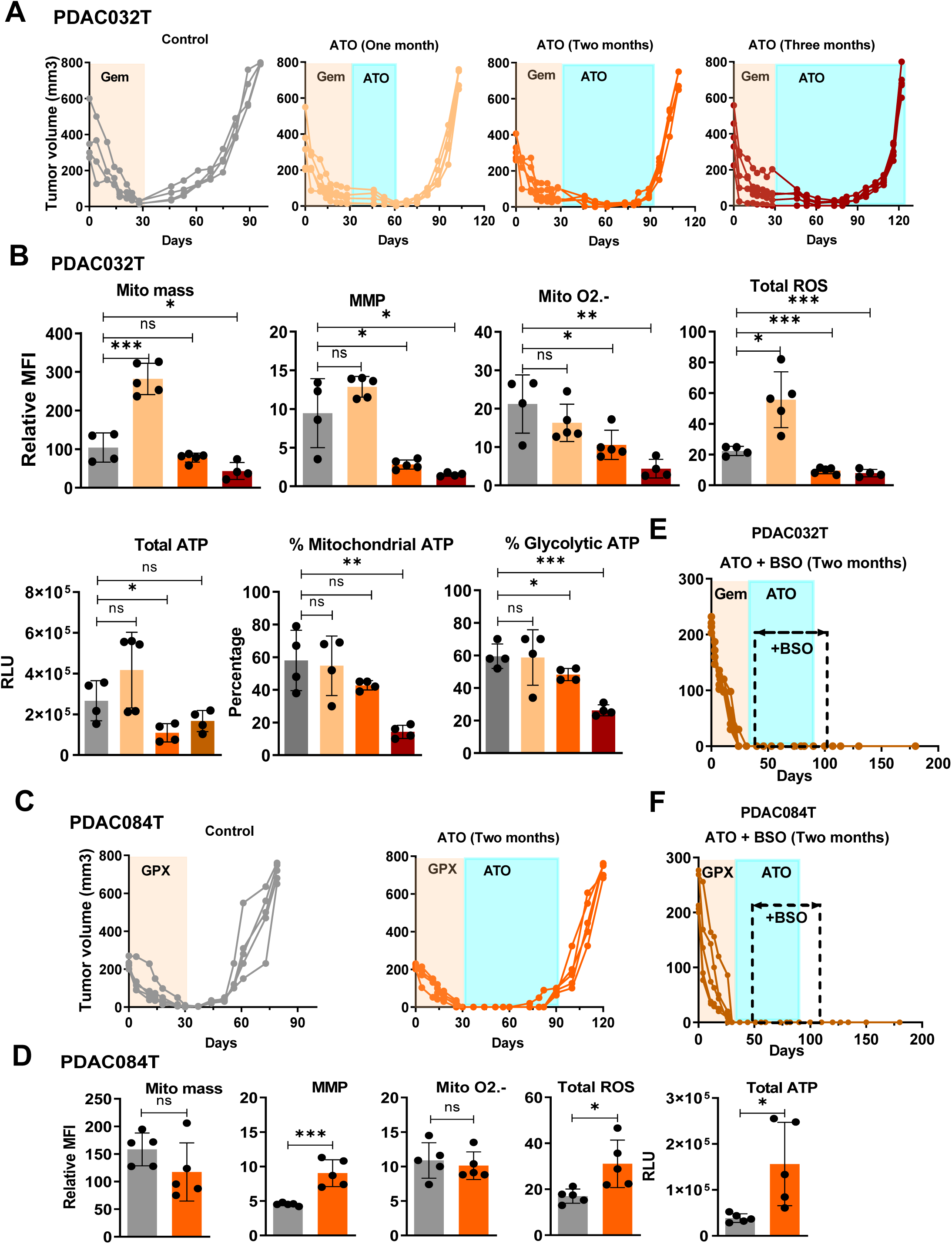
Targeting redox metabolism prevents relapse in both PDAC xenografts. **(A)** Tumor volume in PDAC032T xenograft treated with ATO (0.2mg/kg/day) for one, two or three months starting just after the end of chemotherapy-induced complete regression. **(B)** Mitochondrial mass, MMP, Mitochondrial O2.- and total ROS level were measured in PDAC032T xenografts (Top). Median fluorescence intensity (MFI) is shown relative to unlabeled cells. Total ATP level, percentage of mitochondrial and glycolytic ATP were measured using the Cell viability assay (Cell-Titer Glo Kit) in PDAC032T (Bottom). Specific inhibitors were used: oligomycin (1μM) and 2-DG (100mM) to determine mitochondrial and glycolytic ATP, respectively. **(C)** Tumor volume in PDAC084T xenograft treated with ATO (0.2mg/kg daily) for two months starting after the end of combinatory therapy-induced complete regression. **(D)** Mitochondrial mass, MMP, Mitochondrial O2.- and total ROS level were measured in PDAC084T xenografts. Total ATP level was measured using the Cell viability assay (Cell-Titer Glo Kit). **(E and F)** Tumor volume in PDAC032T (**E**) and PDAC084T (**F**) xenografts, respectively, treated with ATO (0.2 mg/kg daily) and BSO (0.3 mg/kg 3 times per week) during two months after the chemotherapy-induced complete regression. Unpaired T-test was used for statistical analyzes comparing each group with the untreated group. *, ** and *** p<0.05, 0.01, 0.001, respectively; ns: non-significant difference.

In addition, a 2-month ATO treatment was performed in PDAC084T xenograft (Fig. 5C). We observed that ATO treatment delayed relapse compared with control (relapse of non-ATO-treated control mice). Tumor growth started to resume before the end of treatment, suggesting escape from ATO treatment. We analyzed mitochondrial characteristics and ROS level in relapsed tumors, and found increased MMP and ROS level as well as ATP level compared with control (Fig. 5D), without any change of mitochondrial mass and mitochondrial superoxide, suggesting higher energy production.

Next, we explored the possibility of greater efficacy in relapse prevention by combining ATO-induced ROS production with a decrease in antioxidant defenses, based on previous studies suggesting that redox combination therapy is a clinically promising approach in the treatment of advanced solid tumors (37). We chose to combine ATO with buthionine sulfoximine (BSO), an inhibitor of glutathione synthesis. BSO treatment was started slightly later than ATO treatment and was performed for 2 months in both xenografts. Importantly, we observed that the combination of ATO and BSO completely prevented relapse in both PDAC xenografts (Fig. 5E and F). We did not observe any relapse during the three months following the end of the combined treatment (we sacrificed the mice at this time). These data suggest that the combination of ROS overproduction (ATO) and inhibition of antioxidant capacity (BSO) might be a promising strategy to prevent PDAC relapse.

## DISCUSSION

In this work, we demonstrate treatment-acquired resistance in PDAC *in vivo*, through tumor growth from drug-tolerant persister (DTP) cancer cells. These DTP cells survive during treatment due to redox metabolic adaptations (schematic model in Supplementary Fig. S8). The existence of DTP cells in tumors was demonstrated during the last decade and their major biological features were defined: lack of additional genomic alteration, stagnant cell proliferation, reversible drug sensitivity, and flexible energy metabolism (12,13,38,39). These cells are difficult to study due to their low number in the minimal residual disease (scar of the regressed tumor). This is why we analyzed tumors under treatment just before total regression and at the onset of tumor relapse, finding the same metabolic characteristics as in recurrent tumors with bigger volume at the ethical endpoint.

Regarding the reversible drug sensitivity of DTP cells, we have previously illustrated the response to a second cycle of treatment of patient-derived xenografts which relapse after treatment-induced regression (30). With regard to the characteristic of flexible energy metabolism, we show here that DTP PDAC cells exhibit higher mitochondrial oxidative metabolism than untreated PDAC cells (increased mitochondrial mass, active polarized mitochondria, increased ATP production, high ROS, and antioxidant levels). These characteristics have been observed in several different cancers such as melanoma, acute myeloid leukemia, colorectal, and breast cancer (13). Importantly, this work is the first demonstration of the existence of PDAC DTP cells *in vivo*. The experimental background in the pioneer study of Viale et al. in 2014 (26) was that of cells surviving genetic ablation of the Kras oncogene responsible for tumor relapse and relying on mitochondrial respiration to survive, whereas our experimental model implementing drug treatments is a true context of persistence supported by drug tolerance.

New therapies that effectively eradicate resistant cancer cells and residual cells after a therapeutic response are an urgent medical need. A better knowledge of the mechanisms of therapy-induced drug resistance is required to develop strategies to prevent the emergence of drug-resistant cells (34,35,38). Targeting mitochondrial metabolic adaptations, as in chronic myeloid leukemia (17), could not be used in PDAC because these adaptations also occur in tumors treated with gemcitabine combined with perhexiline, which inhibits mitochondrial activity. In contrast, we show that targeting the adaptations in redox metabolism that arise during treatment is a promising therapeutic approach for preventing relapse in PDAC. We have exploited the increased ROS and antioxidant content in DTP cells, which promote cell survival, to implement a strategy aimed at inducing severe oxidative stress likely to promote DTP cell death. This was achieved by the use of drugs that further increase intracellular ROS levels, combining increased ROS production with a pro-oxidant and disruption of the cellular antioxidant system (35,40).

The strength of our study is the demonstration for the first time of PDAC DTP cells *in vivo* (and not *in vitro*), in two different models of PDAC mice, either immunodeficient (heterotopic xenografts) or immunocompetent (orthotopic syngeneic allografts), both responding to chemotherapeutic treatment but relapsing after complete regression. Furthermore, we illustrate for the first time the possibility of preventing tumor growth relapse from DTP cells by inducing their death through strong oxidative stress. This strategy remains to be implemented in immunocompetent preclinical models considering the importance of tumor microenvironment immune cells in the antitumoral response and DTP clearance (13).

Collectively, combining the promotion of ROS production and the inhibition of antioxidant capacity could be a promising avenue for treating pancreatic cancer in the clinic.

## Authors’ Disclosures

The authors declare no conflicts of interests.

## Authors’ Contribution

**N. Abdel Hadi:** Data curation, formal analysis, validation, investigation, visualization, methodology, writing–original draft. **G. Reyes-Castellanos:** Conceptualization, formal analysis, supervision, validation, investigation, visualization, methodology, writing– original draft, writing–review and editing. **T. Gicquel:** Formal analysis, investigation, methodology. **S. Gallardo-Arriaga:** Investigation, writing–original draft. **E. Boet:** formal analysis. **J.-E. Sarry:** formal analysis. **R. Masoud:** Conceptualization, formal analysis, supervision, methodology. **J. Iovanna:** Resources. **A. Carrier:** Conceptualization, resources, data curation, formal analysis, supervision, funding acquisition, validation, visualization, methodology, writing–original draft, project administration.

## Acknowledgments

We thank Nathalie Auphan-Anezin (CIML) for the KPC luc2 cells, the cell culture platform (PCC, TPR2, Marseille, France) for technical assistance, and Karim Sari and Régis Vitestelle for assistance with the use of the PSEA animal housing facility. In addition, we thank Odile and Julie for their advice in cell culture, Martin Bigonnet and Yifan Jiang for their advice on the orthotopic xenografts experiments, Patricia Santofimia-Castaño for antioxidant and mitochondrial gene primers, and Fabio Marchiano for RNA-seq raw data normalization.

This work was supported by Institut National de la Santé et de la Recherche Médicale, Centre National de la Recherche Scientifique, Fondation ARC pour la Recherche sur le Cancer (PJA20181208207). NAH was supported by Association AZM et Saade (Lebanon) and the Fondation pour la Recherche Médicale, GR-C by the CONACYT (Mexico, grant 339091 / 471717) and La Ligue Nationale contre le Cancer (LNCC), SG-A by the CONACYT (Mexico, grant 708538), TG by the French Agence Nationale pour la Recherche (ANR), and RM by the Fondation ARC pour la Recherche sur le Cancer.

## Notes

The authors declare no potential conflicts of interest.

### Competing Interest Statement

The authors have declared no competing interest.

### Summary of Updates

This version of the manuscript has been revised to update the text, since the field of drug-tolerant persister cancer cells has evolved in recent months.

