## Supplementary data: 8 figures for "Adaptation of redox metabolism in drug-tolerant persister cells is a vulnerability to prevent relapse in pancreatic cancer"

### Supplementary Figures: 8

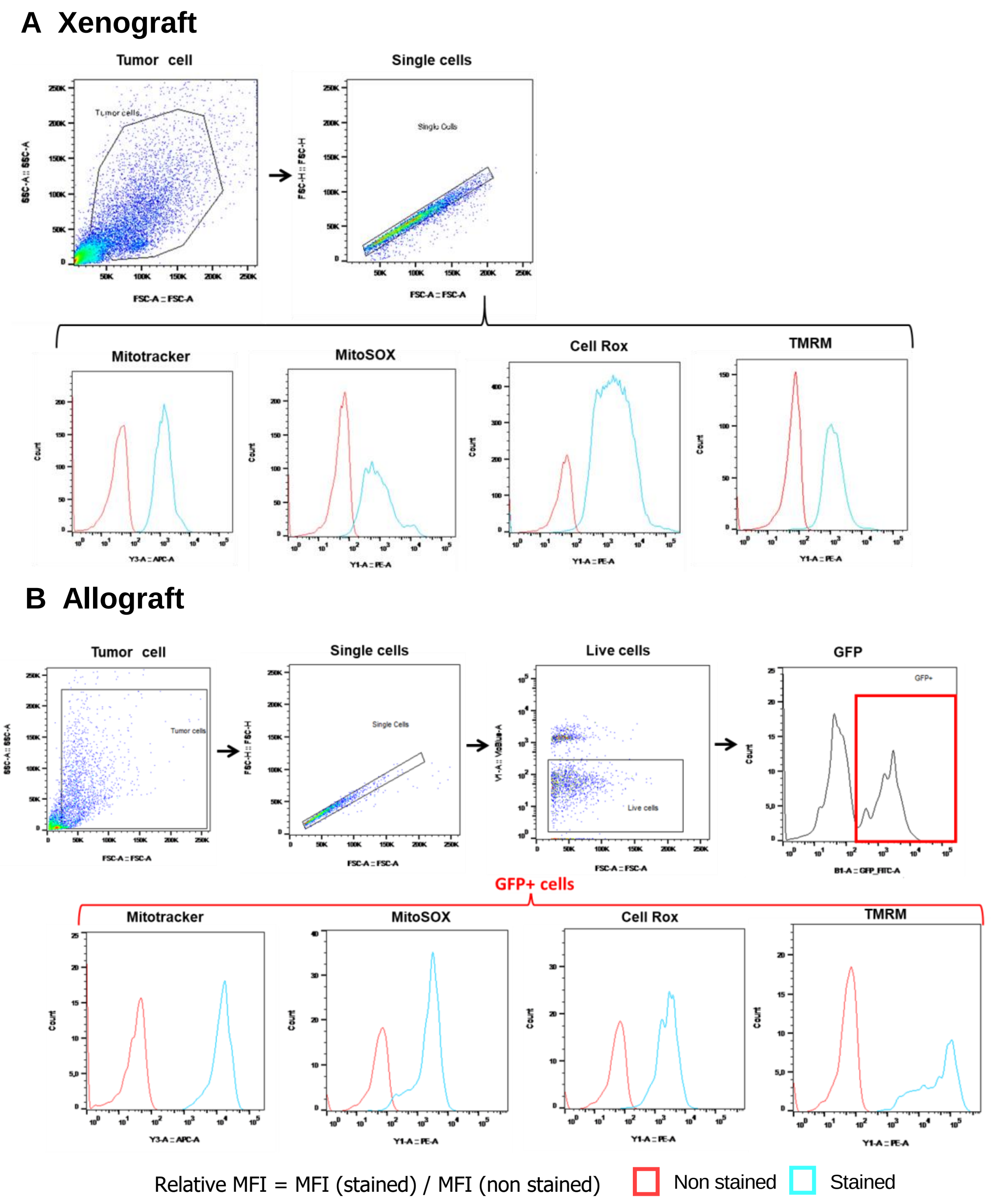


**Figure S1. Representative flow cytometry plots of tumor cells populations from xenograft and allograft mouse models (related to Figures 1 and 2).** Tumor cells are separated from debris by Forward Scatter Area (FSC-A)/Side Scatter Area (SSC-A), and doublets are excluded by FSC-A and FSC-height (FSC-H) dot plots. In xenograft model, Mitotracker deep Red, CellRox Orange, MitoSOX Red, and TMRM histograms are done by gating on single cells **(A).** In allograft model, the histograms are done by gating on GFP$^{+}$ cells (KPCluc2 cancer cells) after gating on live cells (the negative fractions for VioBlue-A staining) **(B).**

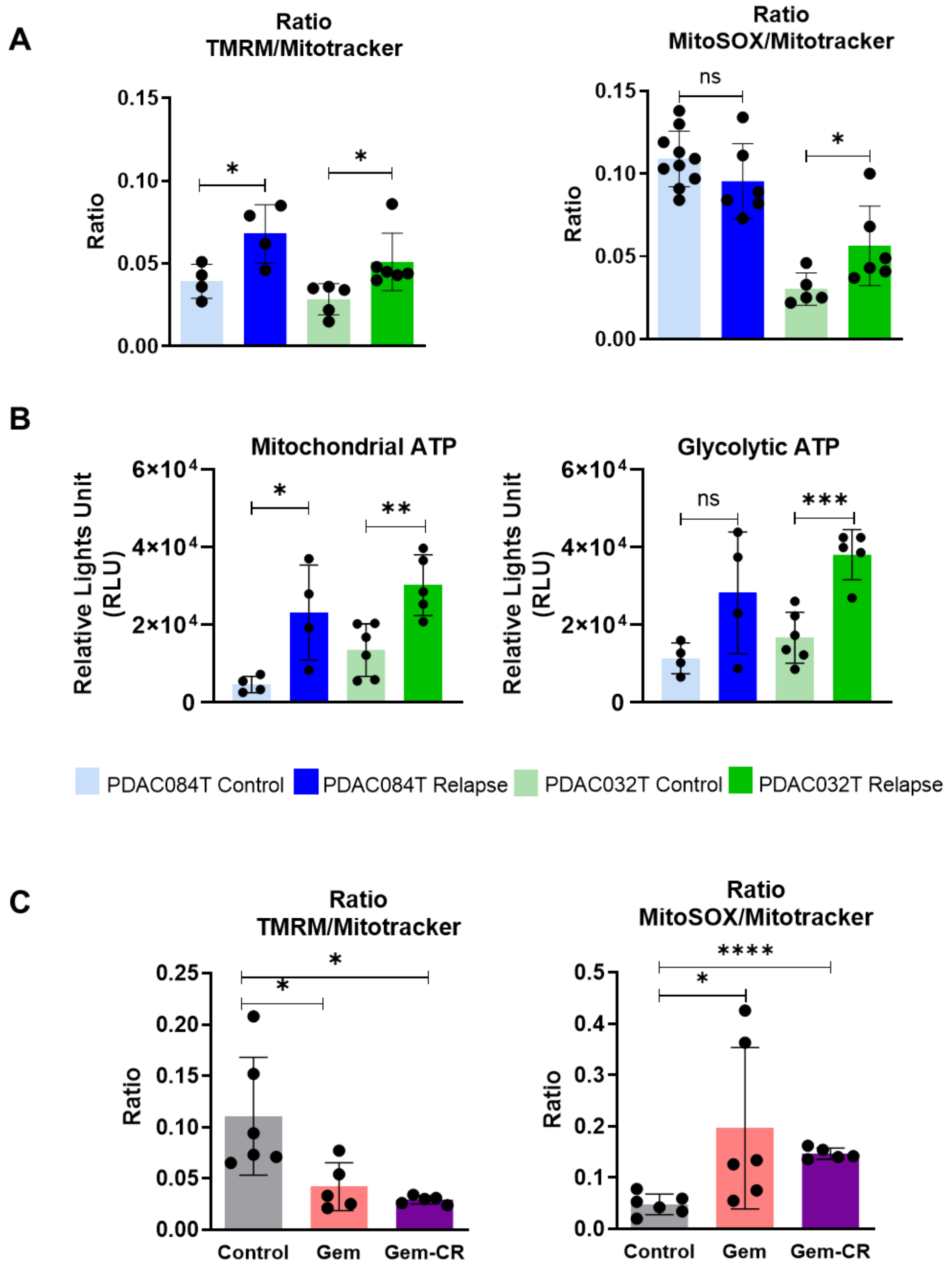


**Figure S2. Mitochondrial and redox metabolic reprogramming imposed by chemotherapy in relapsed PDAC xenografts (related to Figures 1 and 2).** TMRM (measuring mitochondrial membrane potential) and MitoSOX (measuring mitochondrial superoxide anions) values were normalized by Mitotracker (measuring mitochondrial mass) values in relapsed xenografts (**A**), and relapsed allografts (**C**). **(B)** ATP production was measured in the presence of oligomycin (left) and 2DG (right) *in vitro* on dissociated relapsed xenografts, allowing to calculate mitochondrial and glycolytic ATP percentages shown in Figure 1D.

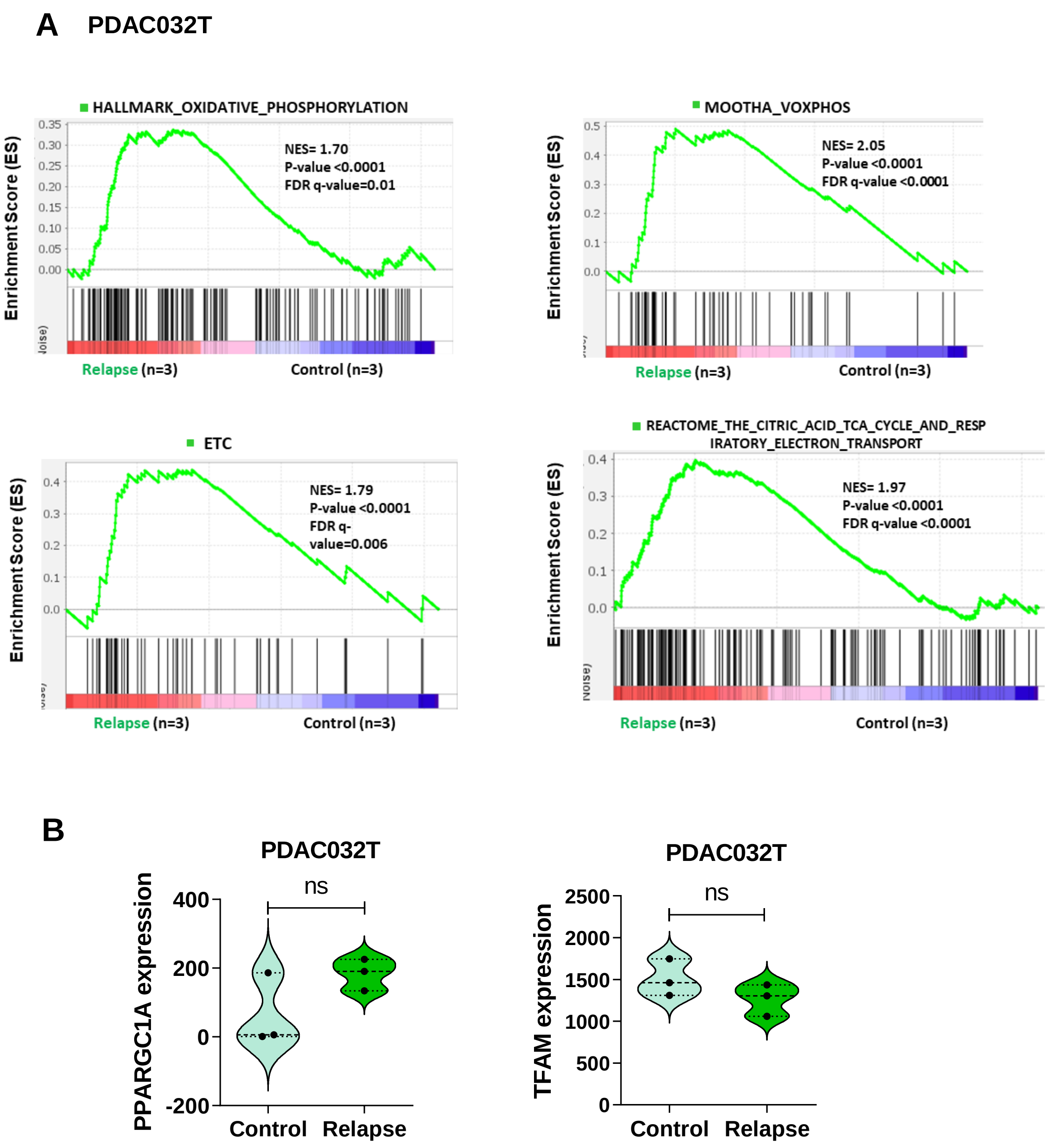


**Figure S3. Gene set enrichment analysis (GSEA) on RNA-seq data from PDAC032T xenografts *ex vivo* (related to Figure 1). (A)** Four mitochondrial signatures were assessed by GSEA: Oxidative phosphorylation, MOOTHA_VOXPHOS, ETC, and Reactome TCA cycle and respiratory electron transport. Significant enrichments are observed in PDAC032T relapsed tumors versus control. The top portion of each panel shows the normalized enrichment scores (NES) for each signature; the bottom portion of the plot indicates the value of the ranking metric moving down the list of ranked genes. **(B)** *PPARGC1A* and *TFAM* expression in PDAC032T relapsed tumors versus control based on RNA sequencing. *P < 0.05; **P < 0.01; ***P < 0.001. ns, not significant.

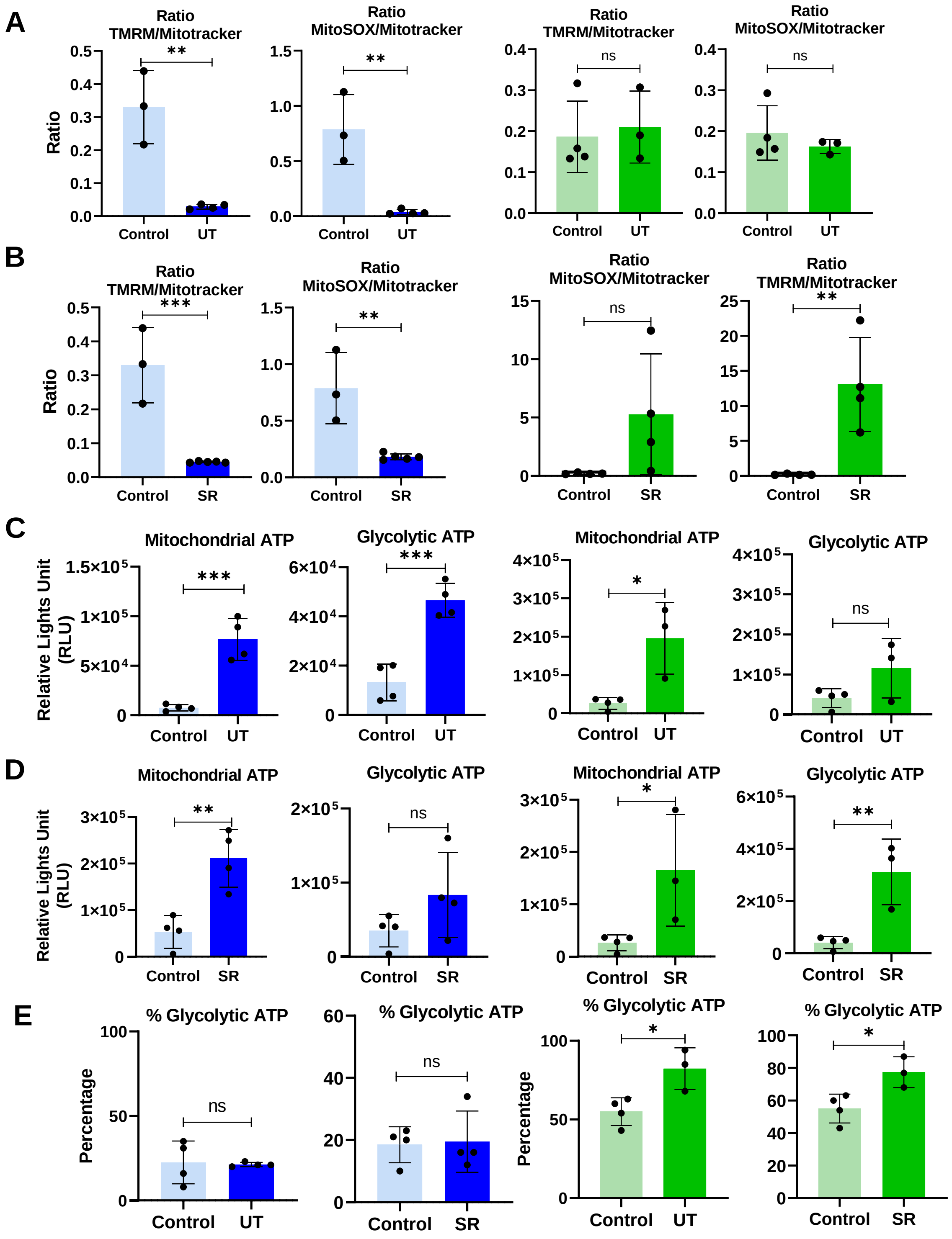


**Figure S4. Mitochondrial and redox metabolic reprogramming occurs during treatment-induced complete regression in PDAC xenografts (related to Figure 3)** **(A-B)** TMRM and MitoSOX values were normalized by Mitotracker values in xenografts PDAC084T (left) and PDAC032T (right), under treatment (UT, **A**) and at start of relapse (SR, **B**). **(C-D)** ATP production was measured in the presence of oligomycin and 2DG *in vitro* on dissociated relapsed xenografts PDAC084T (left) and PDAC032T (right), under treatment (UT, **C**) and at start of relapse (SR, **D**). **(E)** Percentages of glycolytic ATP in PDAC084T (left) and PDAC032T (right), under treatment (UT) and at start of relapse (SR).

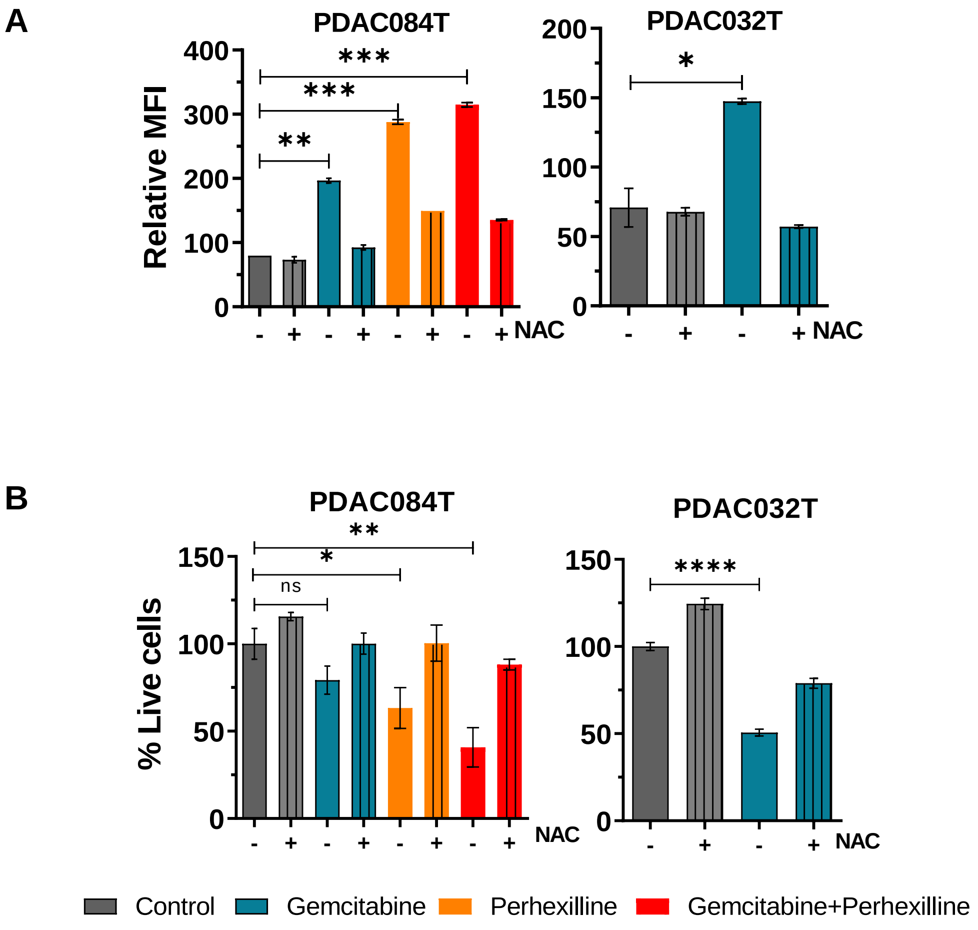


**Figure S5. Chemotherapy induces ROS increase and redox-driven loss of cell viability in PDAC cells *in vitro* (related to Figure 3). (A)** Total ROS level was measured by flow cytometry with the CellROX orange probe in PDAC084T and PDAC032T cells cultured *in vitro*, after 24 hours treatment with gemcitabine (1 µM), or perhexiline (7 µM), or combination at same concentrations. We used DMSO (0.05%) as vehicle for the controls. NAC (2.5 mM) was used as an antioxidant agent to prevent ROS accumulation; supplementation was performed for 24 hours at the same time as the treatments. Data are expressed as mean ± SEM and are representative of three independent experiments performed in duplicates. To calculate p values, Unpaired T-test was used. **(B)** Percentage of live cells treated for 24 hours with gemcitabine (1 µM), perhexiline (7 µM) or the combination in PDAC084T, and with gemcitabine alone in PDAC032T. Cell viability was determined by Crystal violet assay. Data are means of triplicates ± SEM, and are representative of three independent experiments. Unpaired T-test was used. *, **, *** and **** correspond to $p<0.05$, 0.01, 0.001, and 0.0001, respectively; ns: non-significant difference.

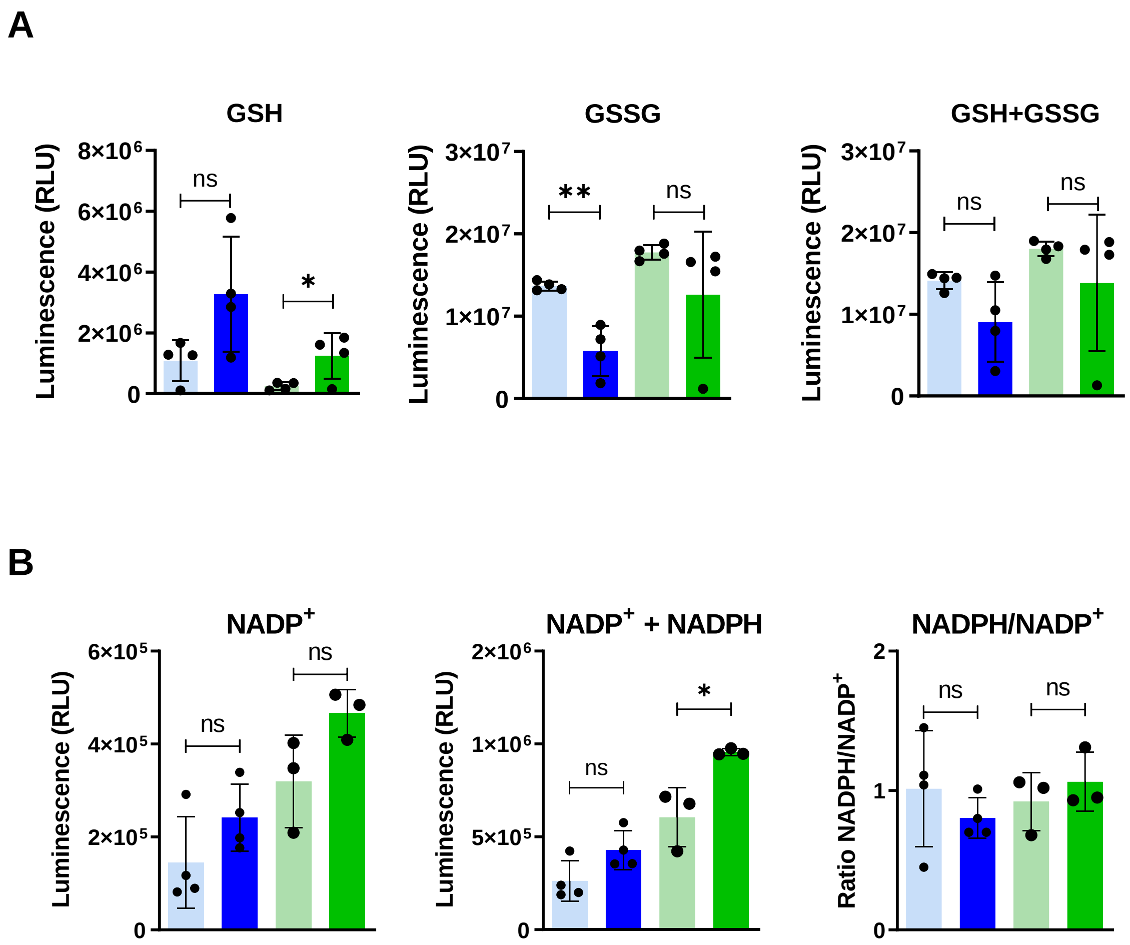


**Figure S6. Abundance of antioxidant small molecules in xenografts at relapse end point (related to Figure 4) (A)** Intracellular levels of GSH, GSSG and total GSH+GSSG levels were measured with the GSH/GSSG-Glo™ Assay| (Promega Kit). **(B)** $NADP^+$ levels, total $NADP^+$ + NADPH levels, and NADPH/$NADP^+$ ratio was measured using the NADP/NADPH-Glo™ Assay kit.

A PDAC032T

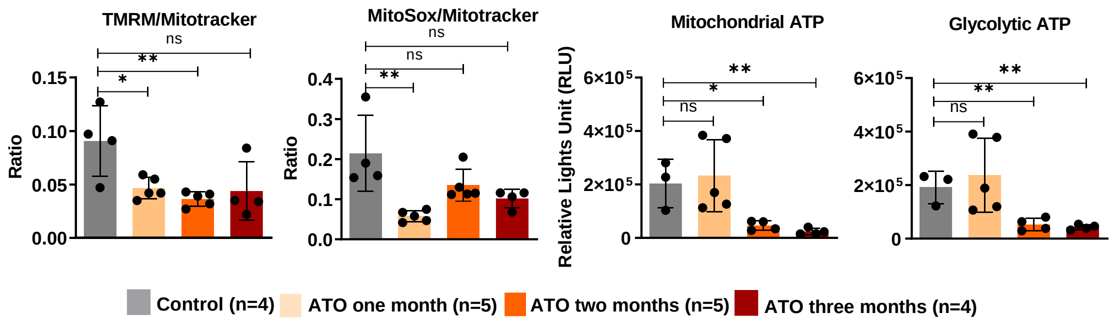


B PDAC084T

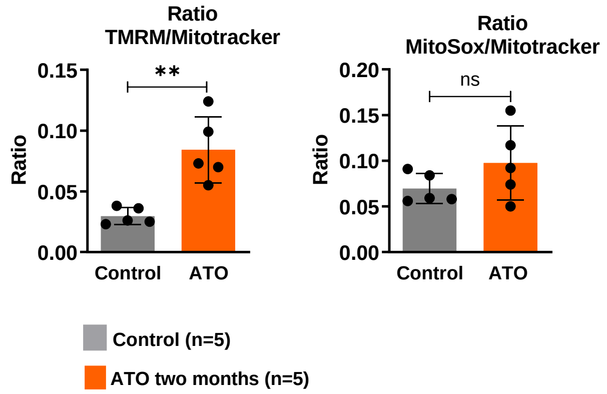


**Figure S7. Mitochondrial metabolic reprogramming during ATO treatment in PDAC xenografts (related to Figure 5) (A-B)** TMRM and MitoSOX values were normalized by Mitotracker values in xenografts PDAC032T (**A, left**) and PDAC084T (**B**), and ATP production was measured in the presence of oligomycin and 2DG in vitro on dissociated xenografts PDAC032T (**A, Right**).

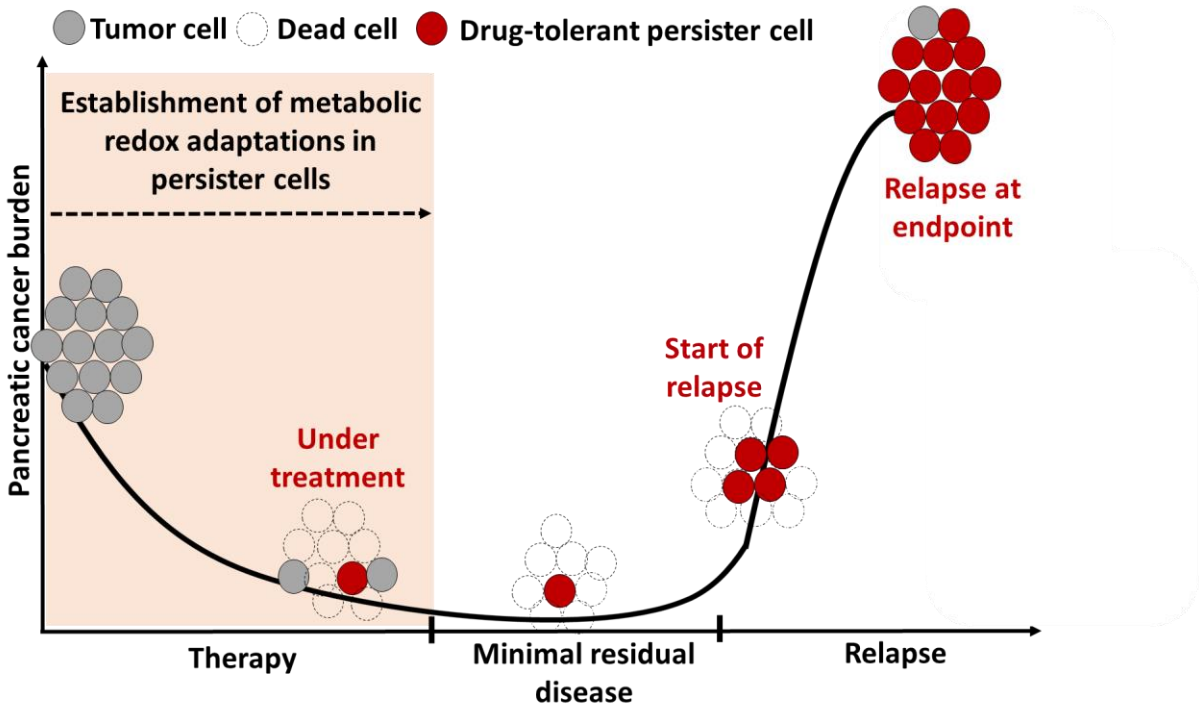


**Figure S8. Working model of treatment-induced acquired resistance in PDAC.** In tumors that respond to therapy, most tumor cells die (dead cells) during treatment, ensuring therapy-induced regression. However, some drug-tolerant persister cancer cells survive during regression, through the establishment of metabolic (mitochondrial and redox) adaptations. They are maintained in the tumor scar and are associated with what is known as minimal residual disease. When these drug-tolerant persister cells resume their proliferation, they are at the origin of the regrowth and the relapse of the tumor.
